# CAR/CXCR5-T cell immunotherapy is safe and potentially efficacious in promoting sustained remission of SIV infection

**DOI:** 10.1101/2021.07.26.453803

**Authors:** Mary S Pampusch, Hadia M Abdelaal, Emily K Cartwright, Jhomary S Molden, Brianna C Davey, Jordan D Sauve, Aaron K Rendahl, Eva G Rakasz, Elizabeth Connick, Edward A Berger, Pamela J Skinner

## Abstract

During chronic human immunodeficiency virus (HIV) or simian immunodeficiency virus (SIV) infection prior to AIDS progression, the vast majority of viral replication is concentrated within B cell follicles of secondary lymphoid tissues. We investigated whether infusion of T cells expressing an SIV-specific chimeric antigen receptor (CAR) and the follicular homing receptor, CXCR5, could successfully kill viral-RNA^+^ cells in targeted lymphoid follicles in SIV-infected rhesus macaques. In this study, CD4 and CD8 T cells from rhesus macaques were genetically modified to express antiviral CAR and CXCR5 moieties (generating CAR/CXCR5-T cells) and autologously infused into a chronically infected animal. At 2 days post-treatment, the CAR/CXCR5-T cells were located primarily in spleen and lymph nodes both inside and outside of lymphoid follicles. Few CAR/CXCR5-T cells were detected in the rectum and lung, and no cells were detected in the bone marrow, liver, brain, or ileum. Within follicles, CAR/CXCR5-T cells were found in direct contact with SIV viral RNA+ cells. We next infused CAR/CXCR5-T cells into ART-suppressed SIV-infected rhesus macaques, in which the animals were released from ART at the time of infusion. These CAR/CXCR5-T cells replicated in vivo within both the extrafollicular and follicular regions of lymph nodes and accumulated within lymphoid follicles. CAR/CXR5-T cell concentrations in follicles peaked during the first week post-infusion but declined to undetectable levels after 2 to 4 weeks. Overall, CAR/CXCR5-T cell-treated animals maintained lower viral loads and follicular viral RNA levels than untreated control animals, and no outstanding adverse reactions were noted. These findings indicate that CAR/CXCR5-T cell treatment is safe and holds promise as a future treatment for the durable remission of HIV.

**Author summary:** A person infected with human immunodeficiency virus (HIV) has replicating virus concentrated within the follicles of lymphoid tissues. The cells needed to clear the infection, cytotoxic T lymphocytes, have limited access to follicles and, thus, the cytotoxic T lymphocytes are never completely able to clear all of the HIV from the body. In this study, we have produced immunotherapeutic T cells that home to follicles and clear infected cells. These T cells express a viral targeting chimeric antigen receptor (CAR) and a molecule called CXCR5, which leads to homing of the cells to follicles. Upon administration of these CAR T-cells to virus-infected primates, we found that the cells localized to the follicle, replicated, and directly interacted with infected cells. While the cells were not maintained in the animals for more than 4 weeks, most of the treated animals maintained lower levels of virus in the blood and follicles than untreated control animals. This study shows that this immunotherapy has potential as a treatment leading to long-term remission of HIV without the need for antiretroviral drugs.

## Introduction

Over 38 million people worldwide live with human immunodeficiency virus (HIV)^1^. Antiretroviral therapy (ART) effectively reduces levels of viral replication in patients; however, ART is not a cure, as it fails to fully eliminate the cellular reservoir of the virus^2–4^. Successful control of HIV-1 requires life-long adherence to ART, which can be challenging for patients with limited or intermittent access to healthcare. Treatment fatigue, side effects^5^, and inconsistent access to medications have led to unsatisfactory levels of treatment adherence that range from 27–80% across various populations, in a disease requiring 95% adherence to be effective. ^6^ Alarmingly, only 57% of people living with HIV (PLWH) in the US are virally suppressed with ART^7^, contributing to the public health threat, as untreated PLWH can transmit virus to others. Increasingly, such transmission can lead to development of drug-resistant strains of HIV^8–10^. To improve the health of individuals with HIV and to reduce community transmission, there is intense global interest in fully eradicating HIV or in developing a strategy for durable remission in the absence of ART^11–17^.

Chimeric antigen receptor (CAR)-T cells have shown great promise in the treatment of certain cancers, including B cell leukemia and B cell lymphomas^18–22^. CAR-T cells are of particular interest in the treatment of HIV, as T cells can be engineered to specifically target the envelope glycoprotein expressed on the surface of HIV-infected cells. In this study, we employed a CAR that targets two independent regions of the viral glycoprotein gp120^12,23^; the CAR contains CD4 (domains 1 and 2) and the carbohydrate recognition domain of mannose-binding lectin (MBL). The MBL prevents CD4 from acting as a viral entry receptor, the bispecific antigen recognition increases the anti-viral potency of the CAR, and the use of self domains minimizes the possibility of immunogenicity^23^.

During chronic HIV and simian immunodeficiency virus (SIV) infections prior to AIDS progression, the vast majority of viral replication concentrates within B cell follicles^24–30^, primarily within T follicular helper cells (Tfh)^24–26,31,32^. Free virions in immune complexes are also localized in follicles through binding to follicular dendritic cells (FDC) in germinal centers (GC)^33–41^. By contrast, levels of virus-specific CD8^+^ T cells, which are critical in controlling HIV and SIV infections, are found at relatively low levels within B cell follicles^26,27,42,43^. In fact, we previously reported the average ratio of in vivo effector (virus-specific CD8^+^ T cells) to target (virus-infected cells) cells is over 40-fold lower in follicular (F) compartments compared to extrafollicular (EF) compartments of secondary lymphoid tissues^27^. Further, we reported that levels of SIV in F areas during early infection^44^ and levels of SIV in both F and EF areas during chronic infection are inversely correlated with levels of virus-specific CD8^+^ T cells in these compartments^27^. In addition, we found that levels of virus-specific CTL in lymphoid tissues correlates with reductions in viral loads ^45,46^. Migration of lymphocytes into B cell follicles is directed by the binding of the chemokine receptor, CXCR5^47–49^, to the chemokine ligand, CXCL13^50–52^, which is produced by follicular stromal cells, such as marginal reticular cells and FDC, and by GC Tfh cells^53–59^. Thus, expression of CXCR5 on the surface of a CD8^+^ T cell mediates migration into B cell follicles^60,61^.

We hypothesize that increasing the numbers of HIV-specific CD8^+^ T cells in B cell follicles will decrease viral replication within the follicles and lead to durable control of HIV^62,63^. In this study, we tested this hypothesis in an SIV-infected rhesus macaque model of HIV infection. Infusion of SIV-infected rhesus macaques (both ART-naïve and ART-suppressed) with rhesus-specific (CD4-MBL)CAR/CXCR5-T cells showed preliminary evidence that the treatment was safe and potentially efficacious in promoting sustained remission of SIV infection. After infusion, CAR/CXCR5-T cells proliferated, accumulated in B cell follicles, interacted with viral (v)RNA^+^ cells, and showed an association with reduced levels of follicular vRNA^+^ cells and overall decrease in plasma viral load. These studies indicate that CAR/CXCR5-T cell immunotherapy shows promise as a tool in the development of durable remission of HIV infection without the need for life-long ART.

## Results

### CAR/CXCR5-T cells home to lymphoid follicles and contact SIV-infected cells in vivo

To evaluate the localization and relative abundance of CAR/CXCR5-T cells in lymphoid and non-lymphoid tissues, and the relative localization within lymphoid tissues of CAR/CXCR5-T cells and SIV vRNA^+^ cells, we autologously infused CAR/CXCR5-transduced T cells into an SIVmac239-chronically infected rhesus macaque and sacrificed the animal 2 days post-treatment (DPT). CAR/CXCR5-T cells were labeled with the fluorescent dye Cell Trace Violet (CTV), and infused into the animal at a dose of 0.35 × 10^8^ cells/kg. Spleen, lymph node (LN), rectum, ileum, bone marrow, lung, liver, and brain were collected at 2 DPT, and examined for localization of the CTV-labeled cells (Figure 1a). CAR/CXCR5-T cells accumulated primarily in the B cell follicles in the F and EF areas of the spleen and LN, with a few cells detected in the rectum and lung. No CAR/CXCR5-T cells were detected in ileum, bone marrow, liver, or brain tissue (Figure 1a). To evaluate the localization of CAR/CXCR5-T cells relative to SIV vRNA^+^ cells, we used duplex RNAScope in situ hybridization (ISH)^64–66^, which allows simultaneous detection of both gammaretroviral vector-transduced CAR/CXCR5-T cells and SIV-infected cells. Spleen tissue sections were hybridized to two sets of probes, one that specifically binds the gammaretroviral CAR/CXCR5 construct and another that specifically binds SIV vRNA (Figure 1b). In addition to detecting SIV vRNA^+^ cells, SIV virions trapped by the follicular dendritic cells (FDC) network were detected as a white haze within the B cell follicle as described previously^60,65^. The duplex RNAScope ISH was combined with immunofluorescence staining to allow the delineation of F and EF areas in lymphoid tissues. CAR/CXCR5-T cells were detected primarily in F areas of the spleen (20.1 cells/mm^2^ in F compared to 3.8 in EF) and, in some instances, were detected in direct contact with SIV vRNA^+^ cells (Figure 1b). We found 4% (23/621) of SIV vRNA^+^ cells were in direct contact with CAR/CXCR5-T cells.

**Fig 1.**
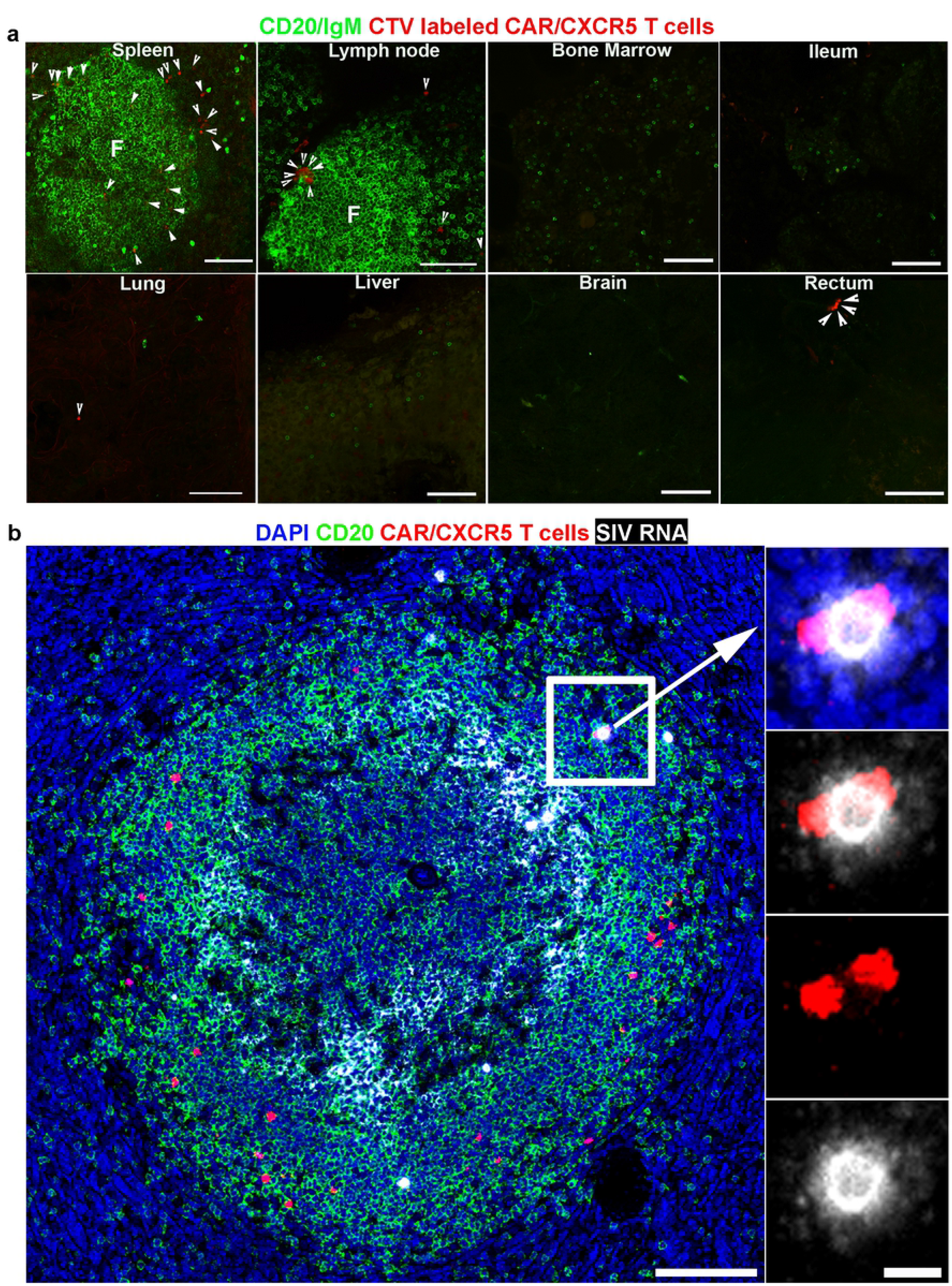
CAR/CXCR5-T cells home to lymphoid follicles and recognize SIV-infected cells in vivo. Location of cell trace violet (CTV)-labeled CAR/CXCR5-T cells was determined in a chronically SIV-infected ART-naïve rhesus macaque, animal R14025 at 2 days post-treatment (DPT). (a) Representative images from spleen, lymph node (LN), bone marrow, ileum, lung, liver, brain, and rectum. Infused cells were labeled with CTV (pseudo-colored red). Tissues were stained with anti-IgM or anti-CD20 (green) to label B cells and delineate B cell follicles (green). Arrowheads point to CTV-labeled cells. Scale bars=100 μm. (b) Representative image of spleen tissue section from animal R14025 showing duplex detection of CAR/CXCR5 construct (red) and SIV (pseudo-colored white) using RNAscope ISH combined with immunofluorescence using a custom-made probe for detection of CAR/CXCR5 construct and a probe specific for SIV. The white haze within the B cell follicle represents SIV virions trapped by the follicular dendritic cells (FDC) network. Scale bar=100 μm. The right panels of panel (b) are enlargements showing an interaction between two CAR/CXCR5-transduced T cells and an SIV-infected cell. The tissue was stained with DAPI (blue), and anti-CD20 (green) to label B cells and delineate B cell follicles. Scale bar=10 μm. Confocal images were collected using a 20× objective. The curves tool in Photoshop was used to increase the contrast of each image in a similar manner.

### Infusion of CAR/CXCR5-T cells into SIV-infected rhesus macaques is safe

We next investigated the safety and in vivo efficacy of CAR/CXCR5-T cell immunotherapy in SIV-infected ART-suppressed animals, compared to untreated control animals. The untreated and treated animal groups included male and female animals, of similar age, weight, peak viral loads and CD4/CD8 frequencies (Table 1).

**Table 1:** Animal information

| Group | Animal | Age (yrs) | Sex | Weight (kg) | Peak VL Post-infection | Pre-ART CD4/CD8 frequency | ART treatment (days) | Necropsy (DPT) |
| --- | --- | --- | --- | --- | --- | --- | --- | --- |
| T0 | R14025 | 3.6 | M | 5.5 | 3.80 x10<sup>7</sup> | - | - | 2 |
| T1 | R01093 | 17 | F | 7.6 | 0.94 x10<sup>7</sup> | 0.44 | 337 | 56 |
| T1 | Rh2526 | 9.8 | F | 5.0 | 4.74 x10<sup>7</sup> | 0.76 | 379 | 70 |
| T1 | Rh2537 | 13 | F | 8.0 | 0.94 x10<sup>7</sup> | 1.28 | 358 | 83 |
| C | R12049 | 6.3 | F | 6.4 | 6.37 x10<sup>7</sup> | 1.06 | 337 | 56 |
| C | R11100 | 7.1 | F | 7.4 | 4.62 x10<sup>7</sup> | 0.31 | 379 | 70 |
| C | R10002 | 9 | M | 11.2 | 2.75 x10<sup>7</sup> | 0.77 | 358 | 83 |
| T2 | Rh2850 | 4.7 | M | 6.3 | 0.93 x10<sup>7</sup> | 1.08 | 104 | 308 |
| T2 | Rh2853 | 7.1 | M | 9.2 | 2.00 x10<sup>7</sup> | 1.29 | 76 | 77 |
| T2 | Rh2858 | 6.4 | M | 10.6 | 2.38 x10<sup>7</sup> | 0.68 | 90 | 328 |

CAR/CXCR5-T cells were produced using gammaretrovirus transduction of peripheral mononuclear cells (PBMCs). For the first treatment group (T1), CAR/CXCR5-T cells were generated using PBMCs collected from rhesus macaques during the chronic stage of infection. For the second treatment group (T2), CAR/CXCR5-T cells were generated from PBMCs collected prior to SIVmac251 infection. T1, T2, and control animals were suppressed with antiretroviral therapy (ART) that was initiated 63–68 days post-infection. Animals were released from ART at the time of CAR/CXCR5-T cell infusion and monitored for at least 60 days, as outlined in the study design shown in Figure 2. Blood and tissue samples were collected over time to monitor infused cells and SIV vRNA.

**Fig 2.**
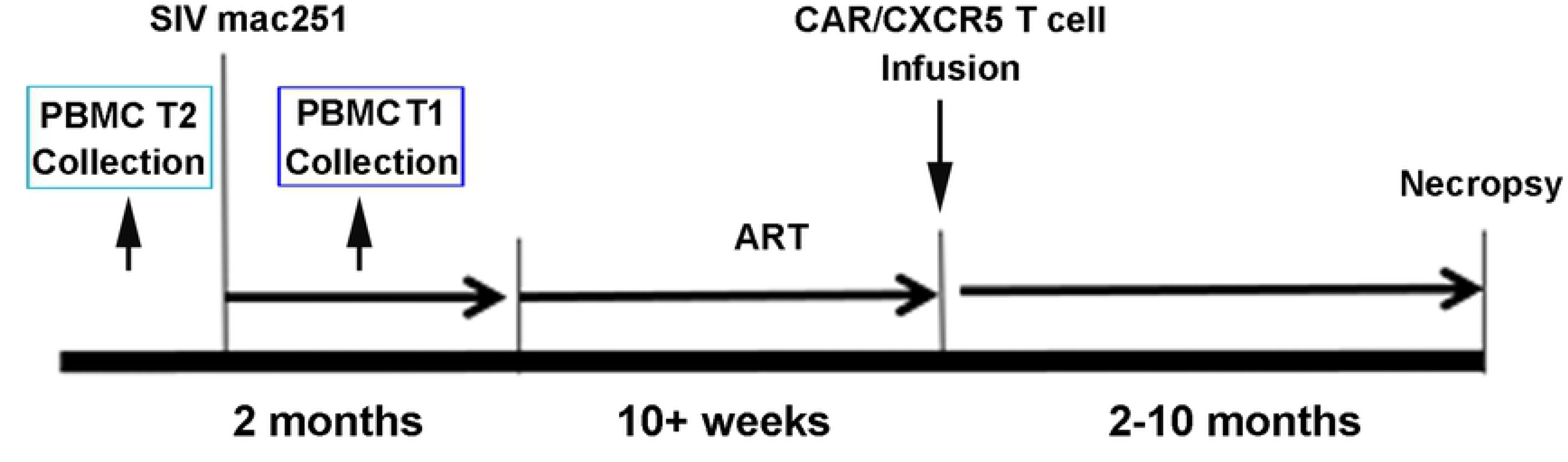
A timeline of the Rhesus macaque pilot studies. Animals were infected with SIV mac251 and ART suppressed. Cells were collected for transduction either post-infection (T1) or pre-infection (T2). ART was interrupted at the time of CAR/CXCR5 cell infusion. Blood and tissue samples were collected at regular intervals post-infusion.

To evaluate the safety of the treatment, animals were monitored by veterinary staff twice daily for any signs of pain, illness, and stress by observing appetite, stool, behavior, and physical condition in response to the infused CAR/CXCR5-T cells. The animals exhibited no observable adverse ill effects after receiving the immunotherapeutic cells, and their weights were unaffected by the immunotherapeutic infusion. Necropsy reports noted no abnormalities in treated animals beyond those typical in SIV-infected animals.

A Luminex assay for monitoring cytokine levels after the cell infusion showed a transient spike in IL-6 and interferon gamma (IFN-γ) at 2 DPT in three of the six treated animals; and levels returned to normal by 6 DPT (Fig S1a–d). Flow cytometry (Fig S1e) and quantitative polymerase chain reaction (qPCR) (Fig S1f) detected cells in bronchoalveolar lavage (BAL) samples shortly after infusion; however, we detected no CAR/CXCR5-T cell accumulation in lung tissues (Figure 1). The cells likely accumulated in the BAL shortly after infusion due to pulmonary circulation. Upon necropsy, the lungs appeared healthy. The overall health of the animals and the transient nature of the cytokine spikes suggests that the infusion of autologous CAR/CXCR5-T cells is safe.

### Infused CAR/CXCR5-T cells were predominantly activated central memory T cells

T1 and T2 animals were infused with CAR/CXCR5-T cells at a dose ranging from 0.8–2.0 × 10^8^ cells/kg (Table 2). The infused cells were a mix of CD8 and CD4 T cells. Most of the infused cells expressed both the CAR and CXCR5 (range, 55–79.4%) and displayed primarily a central memory phenotype (range, 50.3–73.6%) (Table 2, Fig S2). The majority of the central memory cells expressed C-C chemokine receptor 7 (CCR7) (range, 44.6–93.5%), a LN homing molecule^67–69^.

**Table 2:** Cell infusions

| Group | Animal | Cells/kg infused | %CAR/CXCR5 | % CM | % CCR7+ | % CD4<sup>+</sup> | % CD8<sup>+</sup> | % CD4<sup>+</sup>CD8<sup>+</sup> |
| --- | --- | --- | --- | --- | --- | --- | --- | --- |
| T0 | R14025 | $0.35 \times 10^8$ | 66.2 | 58.1 | 93.5 | 43.8 | 16.3 | 38.0 |
| T1 | R01093 | $0.8 \times 10^8$ | 55.0 | 50.3 | 66.5 | 22.3 | 46.5 | 28.8 |
| T1 | Rh2526 | $1.12 \times 10^8$ | 59.5 | 65.3 | 62.5 | 26.8 | 36.0 | 35.4 |
| T1 | Rh2537 | $0.75 \times 10^8$ | 57.4 | 66.9 | 59.0 | 42.3 | 27.3 | 28.9 |
| T2 | Rh2850 | $2.0 \times 10^8$ | 79.4 | 73.6 | 62.5 | 20.6 | 14.3 | 63.8 |
| T2 | Rh2853 | $1.06 \times 10^8$ | 68.6 | 54.5 | 44.6 | 32.3 | 15.3 | 48.3 |
| T2 | Rh2858 | $1.26 \times 10^8$ | 58.4 | 61.4 | 76.4 | 33.4 | 23.2 | 41.9 |

### CAR/CXCR5-T cell infusion associated with reductions in viral loads

Viral loads were monitored after infusion of CAR/CXCR5-T cells into rhesus macaques. Untreated control animals (Figure 3a) showed a rapid rise in viral loads 10 to 14 days after ART cessation, followed by a slow decline over time that remained at detectable levels throughout the 56 to 83 day experiment. T1 animals (Figure 3b), in which cells were collected during chronic untreated infection, showed an initial immediate spike in viral loads due to the presence of virus in the infused SIV-infected transduced cells^70^. Viral loads dropped in all three treated animals— reaching undetectable levels in one of the three animals—and then began to rise. For the duration of the experiment, one of the three T1 treated animals maintained substantially lower viral loads compared to control untreated animals, and two of the three treated animals had undetectable viral loads at necropsy (56-83 DPT).

**Fig 3.**
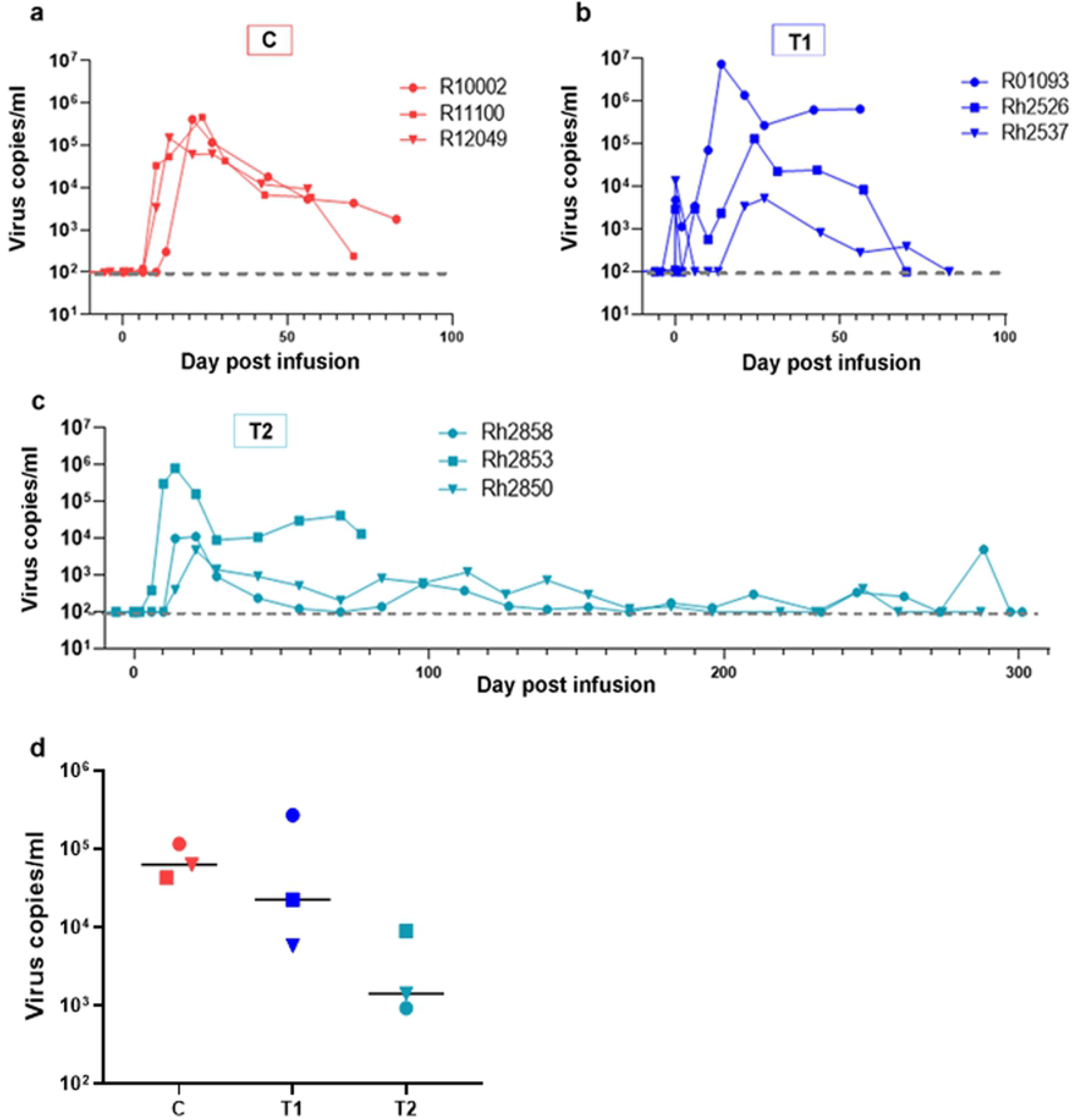
Post-infusion viral loads. Viral loads over time in (a) untreated control animals, (b) pilot study T1 animals and (c) pilot study T2 animals. Viral loads were determined by measurement of gag mRNA relative to the housekeeping gene beta-actin in cell pellets using reverse transcription (RT) polymerase chain reaction (PCR). (d) Viral loads for the three animal groups one month post-infusion (27-30 DPI). The bar represents the median.

In T2 animals (Figure 3c), two of the three treated animals showed lower peak viral loads post-ART release compared to control untreated animals. One-month post-infusion (27–30 DPT), the viral loads in two of the three T1 and all three T2 animals were lower than untreated control animals, with the median viral load of T2 being nearly 2 logs below that of the untreated control animals (Figure 3d). As an exception to the study design, which was planned to maintain animals for only 2–3 months post-cell infusion and ART release, we maintained two T2 animals for 10 months post-infusion to monitor long-term viral loads. During this time, the animals maintained control of infection, with viral loads oscillating between undetectable and very low levels (Figure 3c). We detected similar levels of naturally occurring SIV-specific (Mamu-A1*001/Gag-CM9) CD8^+^ T cells in PBMCs of treated and untreated control animals at one-month post-infusion (Fig S3). This finding suggests that the differences in viral loads between groups was not likely driven by differences in the endogenous response. Overall, these data suggest that CAR/CXCR5-T cell therapy is effective at reducing viral loads in SIV-infected rhesus macaques after ART cessation.

### CAR/CXCR5-T cells proliferate in vivo

At 2 DPT, CTV-labeled CAR/CXCR5-transduced cells showed evidence of proliferation in both the EF and F areas of LNs, showing doublets of cells and cells with decreased fluorescence intensity, indicating a loss of CTV with cell division. Cells within the F areas had an overall lower CTV fluorescence intensity than those in the EF (Figure 4a), suggesting that they had undergone further cell division. Similarly, at 2 DPT, RNAScope detection of CAR/CXCR5-T cells combined with immunofluorescence staining of LNs showed clusters of CAR/CXCR5-T cells at the edge of the follicles, suggestive of cell expansion (Figure 4b). These clusters were detected in over 50% of the follicles (range, 49–58%). In vivo proliferation of CAR/CXCR5-T cells was further confirmed at 6 DPT in LN sections in treated animals by a combination of RNAScope and Ki67 antibody staining to mark T cell activation and proliferation. We detected Ki67^+^ CAR/CXCR5-T cells in both F and EF areas (Figure 4c). Levels of Ki67^+^ CAR/CXCR5-T cells showed a median of 30% (range, 9–64%) of total CAR/CXCR5-T cells in F areas and a median of 36% (range, 13-44%) of total CAR/CXCR5-T cells in EF areas. Interestingly, the T2 animal demonstrating the greatest control (Rh2850), showed the highest percentage of F Ki67^+^ CAR/CXCR5-T cells (64%) and the animal that lost control (Rh2853) showed the lowest percentage of follicular Ki67^+^ CAR/CXCR5-T cells (9%) (Figure 4d).

**Fig 4.**
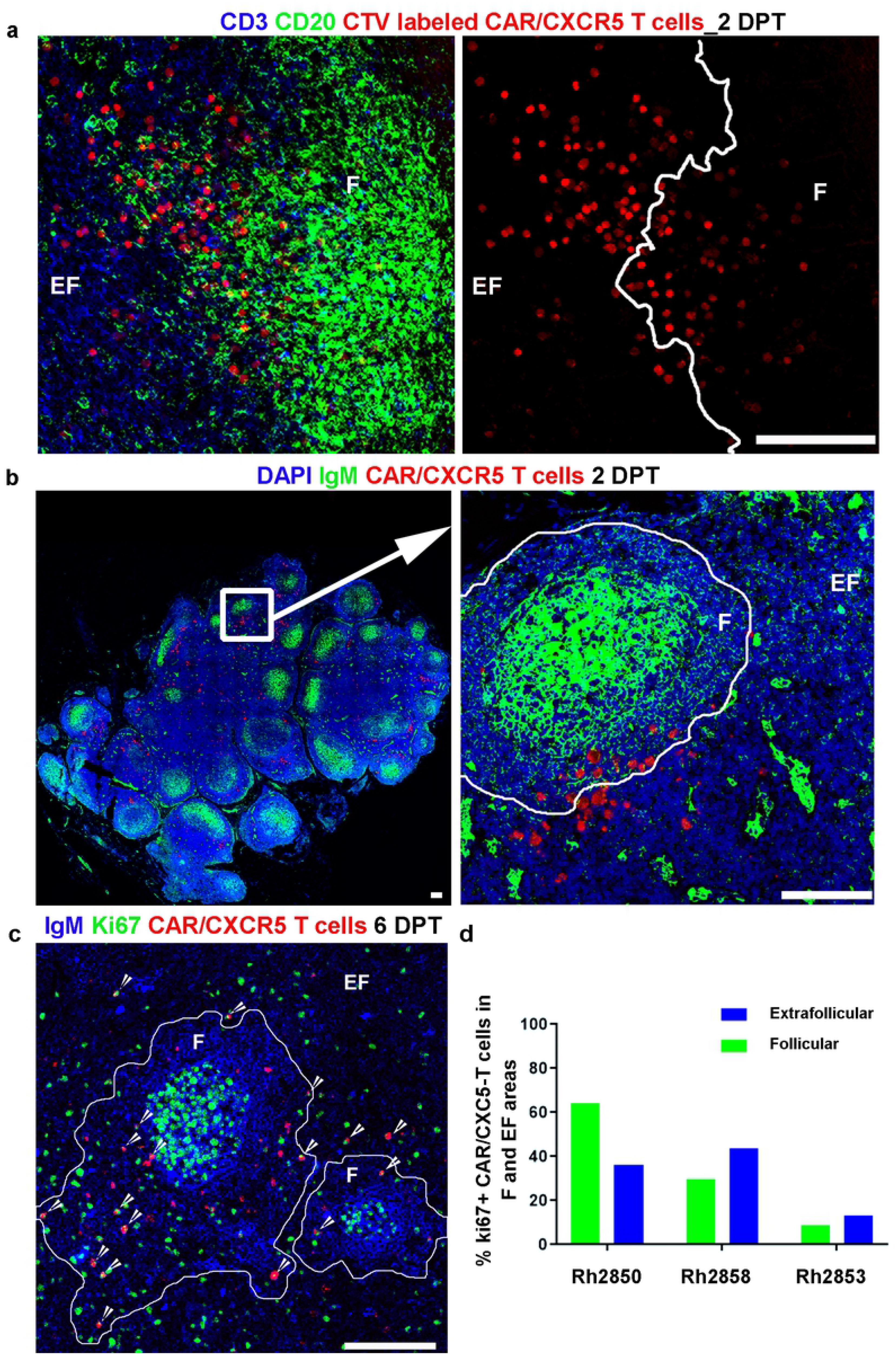
CAR/CXCR5-T cells continue to expand in vivo. CAR/CXCR5-T cells expanded at the edges of B cell follicles in vivo at 2 DPT and accumulated within B cell follicles at 6 DPT. (a) Representative image from LN tissue from Rh2850 stained 2 DPT showing CAR/CXCR5-T cell proliferation in the extrafollicular (EF) area near the follicular (F) zone. Tissues were stained with anti-CD3 (blue) to label T cells and anti-CD20 (green) to label B cells and delineate B cell follicles (F). CAR/CXCR5-infused cells were labeled with CTV (pseudo-colored red). Cells within F showed lower fluorescence intensity indicating division. Scale bar=50 μm. (b) Representative image of LN tissue section, from Rh2858 visualized using RNAScope ISH combined with IF, showing expansion of CAR/CXCR5-T cells at the edge of follicles at 2 DPT. The right panel is an enlargement from left panel showing a cluster of CAR/CXCR5-T cells (red) that appear to be expanding at the edge of the follicle. Tissues were stained with DAPI (blue) and anti-IgM (green) to label B cells and delineate B cell follicles (F), with the more brightly stained germinal center in the center of the F. (c) Representative image of LN tissue from Rh2850, showing CAR/CXCR5-T cell (red) proliferation in F and EF at 6 DPT detected by RNAScope ISH combined with IF. Tissues were stained with anti-IgM (Blue) and anti-Ki67 (green) to mark activation and proliferation. B cell follicles are delineated with white lines. Scale bars=100 μm. Confocal images were collected using a 20× objective. (d) Percentage of Ki67+ CAR/CXCR5 T cells in the F (green) and EF (blue) areas for each of the T2 animals.

### CAR/CXCR5-T cells localize to the follicle and persist for up to 28 days

We analyzed CAR/CXCR5-T cells in LN sections from T2 animals biopsied at 2, 6, 14, 28, and 60 DPT using RNAscope. There was a noticeable shift at 6 DPT to CAR/CXCR5-T cells primarily accumulating within B cell follicles (Figure 5b compared to Figure 4b). Quantification of CAR/CXCR5-T cells in the F and EF regions of LNs revealed that CAR/CXCR5-T cells were most abundant during the first week post-infusion, followed by a decline over time (Figure 5c). At 2 DPT, CAR/CXCR5-T cells were detected at similar levels in both F and EF areas, with a median of 28 cells/mm^2^ (range, 26–30 cells/mm^2^) in F areas and 30 cells/mm^2^ (range, 23–37 cells/mm^2^) in EF areas (Figure 5c). At 6 DPT the cells were detected predominantly in F areas with a median of 78 cells/mm^2^ (range, 34–239 cells/mm^2^) compared to a median of 11 cells/mm^2^ (range, 3–38 cells/mm^2^) in EF areas. (Figure 5c). At 14 DPT, the F:EF ratio increased, however, the overall frequency of cells sharply declined in all treated animals, with a median of 2.3 cells/mm^2^ (range, 0.32–7.6 cells/mm^2^) in F and 0.4 cells/mm^2^ (range, 0–0.57 cells/mm^2^) in EF areas (Figure 5c). By 28 DPT, cells were only detected in F areas of one animal (Rh2850; 1.17 cells/mm^2^), with no cells detected at 60 DPT in any of the examined sections of the treated animals (Figure 5c). Notably, the animal that lost viral control (Rh2853) showed the fastest and steepest decline in levels of CAR/CXCR5-T cells over time relative to two animals that controlled infection.

**Fig 5.**
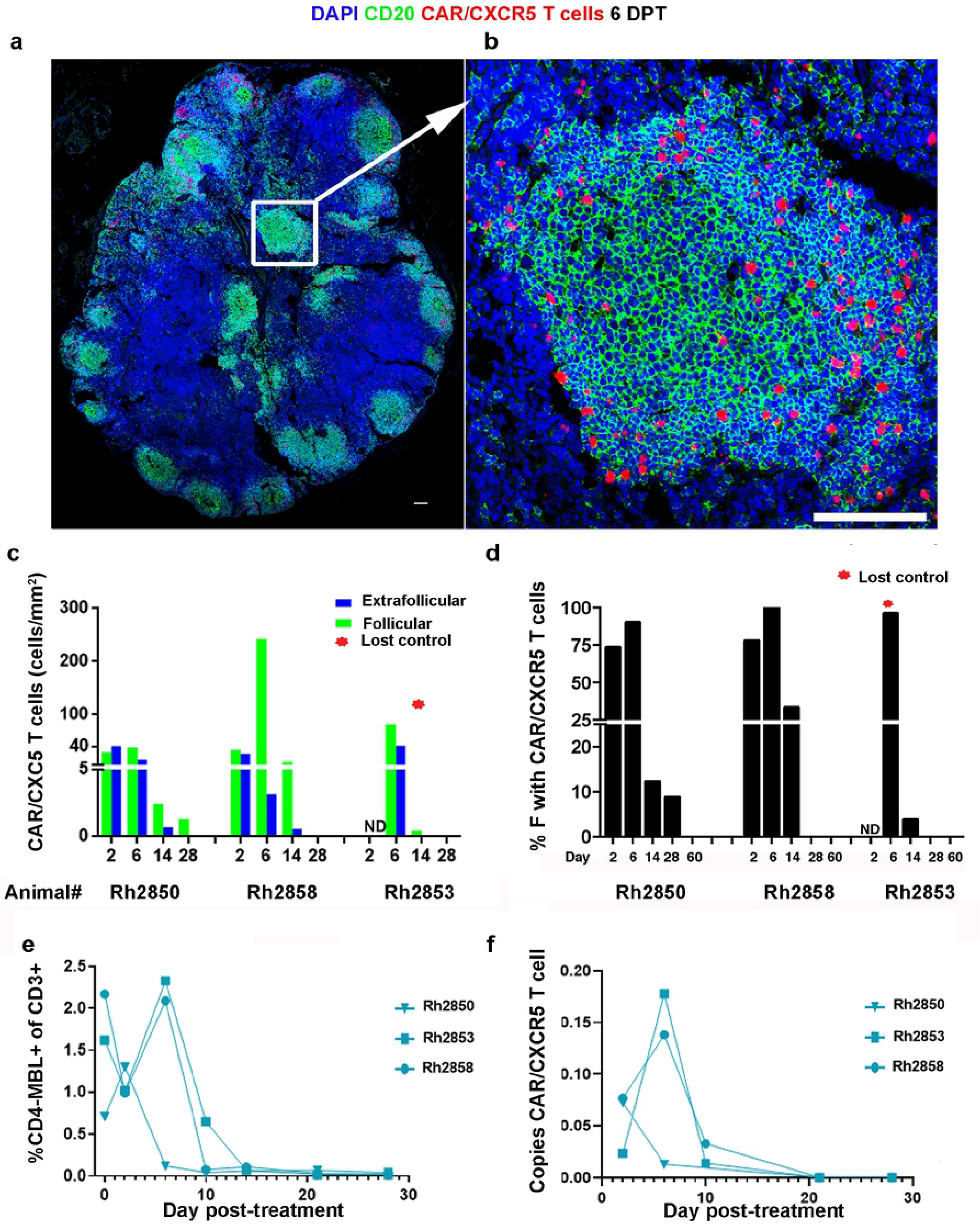
CAR/CXCR5-T cells localize to over 90% of the follicles and persist for up to 28 days. CAR/CXCR5-T cells successfully homed to over 90% of the B cell follicles at 6 DPT and persisted for up to 28 DPT in SIV-infected ART-suppressed/released animals. Representative images of LN tissue section from Rh2858, showing CAR/CXCR5-T cells (red) detected using RNAScope ISH combined with IF using a custom-made probe for detection of the CAR/CXCR5 construct. (a) Confocal image showing a whole LN tissue section. (b) Enlargement of the delineated area in (5a) showing that CAR/CXCR5-transduced T cells (red) successfully homed to a B cell follicle (green). The tissue was stained with DAPI (blue), and anti-CD20 (green) to label B cells and delineate B cell follicles. The confocal image is collected with a 20× objective. Scale bar=100 μm. (c) Levels of CAR/CXCR5-T cells over time after infusion in F (green) and EF areas (blue) of LN. The animal that lost control of the virus infection is marked with a red star. Samples not determined are marked ND. (d) Percentage of follicles that contained CAR/CXCR5-T cells over time post-infusion. (e) Frequency of CD4-MBL^+^ cells in CD3^+^ PBMCs as determined by flow cytometry and (f) copies of CAR in the total cell population in BAL as determined by quantitative real-time PCR at the indicated time points post-infusion.

Most follicles in the LNs had detectable CAR/CXCR5-T cells during the first week post-treatment. In fact, at 6 DPT; a median of 96% (range, 90–100%) of follicles examined had CAR/CXCR5-T cells (Figure 5d). These levels declined in all animals at subsequent timepoints.

Examination of PBMCs using both flow cytometry and qPCR revealed a similar pattern of CAR/CXCR5-T cell persistence. Flow cytometry analysis detected CAR/CXCR5-T in isolated PBMCs up to 14–21 DPT (Figure 5e), and genomic DNA PCR detection of CAR/CXCR5-T cells in PBMCs showed a similar decline in cell number by day 14 (Figure 5f). In addition, we found a strong positive correlation between levels of follicular CAR/CXCR5-T cells in LN tissue in situ and the frequency of CD4-MBL^+^ CAR cells detected in PBMCs by flow cytometry (Fig S4). This finding suggests that the cells have similar persistence in peripheral blood and tissue, and that detection of CAR/CXCR5-T cells in peripheral blood may be a useful surrogate marker for levels and persistence of CAR/CXCR5-T cells in lymphoid tissues.

### In vivo levels of viral RNA appear to be impacted by CAR/CXCR5-T cells

We determined the levels of vRNA in the three T2 animals and three control animals at 28 DPT (Figure 6). The two treated animals that exhibited sustained control of SIV infection (Rh2850, Rh2858) showed few to no SIV vRNA^+^ cells at 28 DPT in F and EF areas compared to abundant SIV vRNA^+^ cells in untreated control animals and the T2 animal that did not control the infection (Rh2853) (Figure 6a and b). Notably, we detected no CAR/CXCR5-T cells that were SIV vRNA^+^ in the examined sections. In addition, Rh2850 and Rh2858 animals had lower percentages of follicles with free virions trapped by the FDC network than untreated control animals, or the treated animal that lost control (Rh2853) (Figure 6c). In fact, Rh2850 had no detectible FDC associated virions in any follicles, and only one of 30 follicles showed FDC-associated virions in Rh2858, whereas most follicles showed FDC trapped virions in untreated control animals and the treated animal that lost control. These findings suggest that the immunotherapeutic cells may have led to sustained reductions in vRNA in the treated animals.

**Fig 6.**
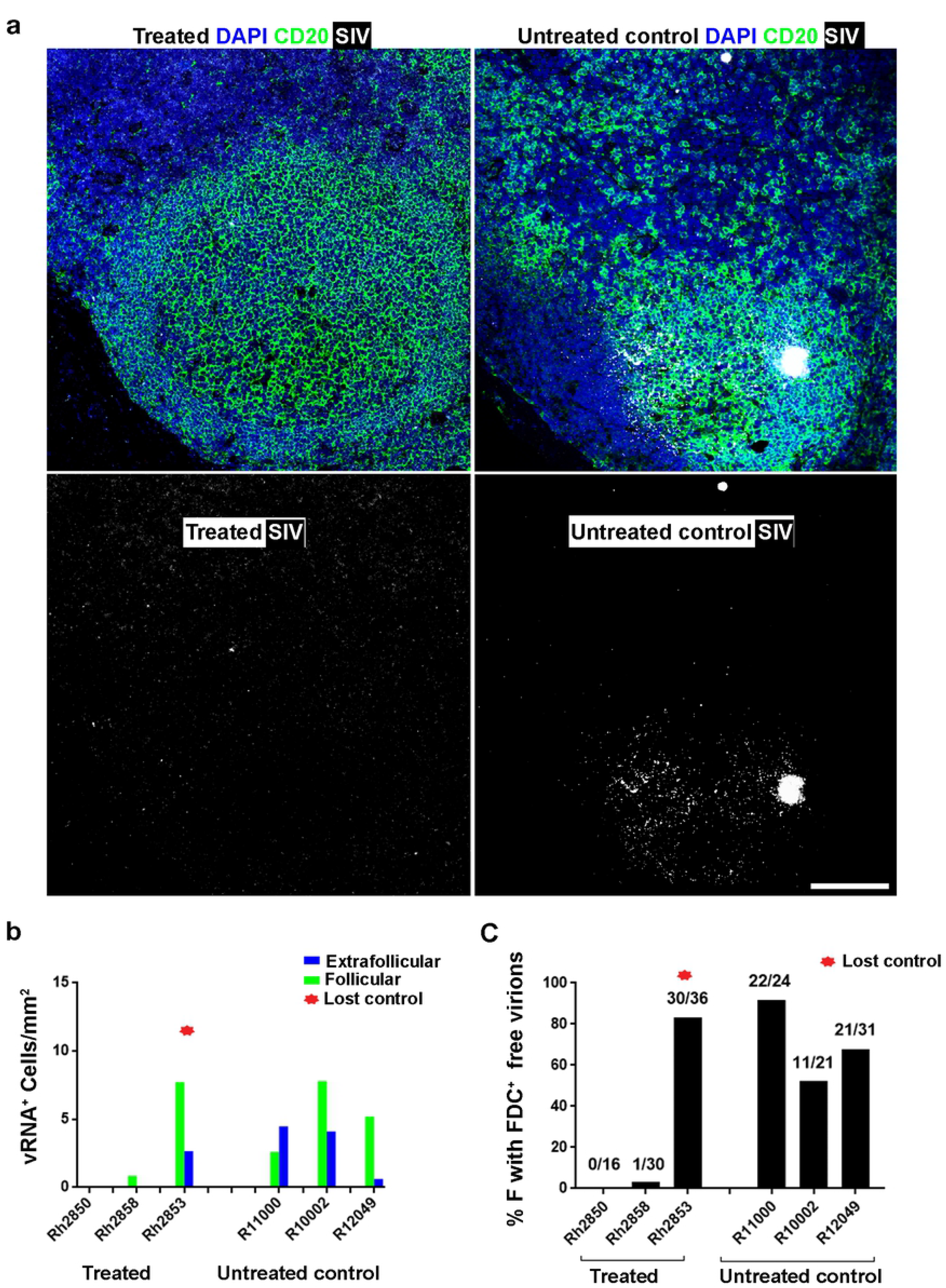
Infusion of CAR/CXCR5-T cells into SIV-infected rhesus macaques results in lower viral loads post-ART interruption. At 28 DPT, treated animals showed a reduction in viral (v)RNA compared to untreated control animals. (a) A representative image from a LN tissue section showing the abundance of SIV vRNA^+^ cells and free virions trapped by follicular dendritic cells network (pseudo-colored white) detected using RNAscope ISH combined with IF in treated from Rh2850 (left panels) versus untreated control from R11002 (right panels). The tissue was stained with DAPI (blue), and anti-CD20 (green) to label B cells and delineate B cell follicles. The white haze within the B cell follicle represents SIV virions trapped by the follicular dendritic cells (FDC) network. Confocal images were collected with a 20× objective. Scale bar =100 μm. (b) Levels of viral RNA in F and EF areas. The animal that lost viral control of the virus infection is marked with a red star. (c) The percentage of follicles with free virions bound by FDC.

## Discussion

HIV and SIV viral replication is concentrated in lymphoid B cell follicles^24–30^ during chronic infection, with infected Tfh in follicles representing a major barrier to HIV eradication^24–27,31,32,63^. The failure of virus-specific CD8^+^ T cells to accumulate in large numbers within the B cell follicles of HIV-infected individuals and SIV-infected rhesus macaques appears to be a major mechanism allowing for persistent follicular viral replication^26,27,32,42,43,71^. In addition, increasing levels of SIV- and HIV-specific T cells within B cell follicles are associated with viral control^27,72–74^. These findings led us to hypothesize that infusion of T cells engineered to co-express a potent SIV-specific CAR along with the B cell homing molecule, CXCR5, will control SIV-infection by reducing viral replication in follicles. We tested this hypothesis in an SIV-infected rhesus macaque model of HIV, in which SIVmac251-infected animals were ART suppressed prior to treatment, and ART removed on the day of treatment with CAR/CXCR5-T cells. Combining a potent SIV-specific CAR with the B cell follicle homing properties of CXCR5 on T cells can overcome the limitation of the endogenous virus-specific CD8^+^T-cell immune response to SIV and lead to control of viremia after ART interruption.

This study provided preliminary evidence of both the efficacy and the safety of autologous CAR/CXCR5-T cell immunotherapy. CAR/CXCR5-T cells successfully homed to B cell follicles and interacted with SIV-infected cells in vivo. The cells proliferated in situ and accumulated primarily in F areas. Five of six treated animals showed lower viral loads at one-month post-infusion and had fewer follicular vRNA^+^ cells in LNs compared to untreated animals. Apart from a transient increase in inflammatory cytokines, none of the animals infused with CAR/CXCR5-immunotherapeutic T cells had an adverse reaction to the infusion. Taken together, these pilot studies in the rhesus macaque model of HIV suggest that autologous CAR/CXCR5-T cell therapy may be a viable tool for the treatment of HIV infection in ART-suppressed individuals after ART cessation.

This study used cells collected either during chronic infection (T1) or collected prior to infection (T2) to produce CAR/CXCR5-T cells. We did not use cells from ART-suppressed animals because we and others observed reductions in the transduction efficiency of T cells from ART-treated subjects^70,75^. Such decreased transduction efficiency may be due to interference with reverse transcription and integration of the gammaretroviral vector by the residual ART drugs. Expression of the transgenes in T cells from ART-suppressed subjects may be improved with alterations in the transduction protocol, use of vectors for virus production that are not impacted by ART drugs or through the use of alternative methods of gene transfer^70^.

Our previous studies examining the function of (CD4-MBL)CAR/CXCR5-T cells indicated that the CXCR5 molecule facilitated migration to B cell follicles both in vitro and ex vivo^61^. Ayala et al. previously showed that CXCR5 expression on T cells induces cell migration into lymphoid follicles in vivo in rhesus macaques^60^. This study extends those findings, demonstrating that the addition of an anti-viral CAR on CXCR5-transduced T cells leads to similar migration and accumulation within B cell follicles.

Shortly after infusion, CAR/CXCR5-T cells proliferated in vivo. At 2 DPT, we detected clusters of replicating CAR/CXCR5-T cells often located at the edge of the follicles. The CAR/CXCR5-T cells appeared to replicate initially next to follicles, then enter into the follicles as suggested by progressively decreasing CTV staining from EF to F areas. Ki67 staining confirmed the in vivo proliferation of CAR/CXCR5-T cells and showed that the cells were replicating in both EF and F areas. It is unclear what cues instigated the in vivo proliferation. Factors that may have contributed include: the culturing conditions prior to infusion, the detection of antigen in vivo by CAR/CXCR5-T cells, or other proliferation cues. To our knowledge, this study is the first to visually confirm the in vivo expansion of autologous SIV-specific CAR T cells in ART-suppressed and -released animals without adding exogenous antigen.

We have previously shown that CAR/CXCR5-T cells readily target and suppress SIV-infected T cells in vitro^61^. This in vivo study showed detection of (CD4-MBL)CAR/CXCR5-T cells directly interacting with SIV-infected cells within lymphoid follicles. These interactions provide evidence that CAR/CXCR5-T cells specifically targeted SIV vRNA^+^ cells, and likely led to the death of these infected cells. Since the CAR does not require antigen presented in the context of an MHC molecule, it is possible that CAR/CXCR5 T cells may not only attack HIV/SIV producing T cells in vivo but may also act to clear the FDCs containing bound SIV. However, it was recently reported that CD4-MBL CAR-T cells did not target FDCs bearing HIV bound immune complexes in vitro^76^, suggesting that the CAR-T cells may not target FDC bearing virus in vivo. We noted little or no virus particles trapped by FDC in the LNs of the T2 animals that went on to show long-term control of the virus and cannot rule out the possibility that the CAR/CXCR5-T cells were able to recognize and remove FDC bearing virions. Alternatively, and perhaps more plausibly, the killing of SIV-producing cells in follicles may have led to a reduction in the seeding of FDCs with virions, explaining the absence of SIV loaded FDC.

Historically, engineering CD8^+^ T cells to express CD4 presents a challenge for HIV treatment due to the likelihood that enhanced CD4 expression would make the transduced CD8^+^ T cells susceptible to HIV infection^12,77–80^ owing to abundant expression of the CCR5 coreceptor^81^. To overcome this issue, we used a bispecific CAR construct, (CD4-MBL)CAR/CXCR5^12,23^. The MBL moiety of the CAR creates a steric hindrance that prevents the CD4 of the CAR from acting as an entry receptor^23^. Nonetheless, CD4+ T cells transduced with CAR and CXCR5 are presumably susceptible to SIV/HIV infections. Interestingly, we detected no SIV vRNA in CAR/CXCR5 cells in the examined sections. These data suggest that either the CD4+ CAR/CXCR5-T cells were not readily infected or, if infected, were rapidly cleared.

Infusion of CAR/CXCR5-T cells appeared to cause no ill effects in the animals, as indicated by veterinary health records and by unremarkable necropsy reports including a lack of inflammation in the lung. Three of six treated animals showed an increase in serum IL-6 and IFN-γ at 2 DPT; however, the effect was transient, did not manifest as detectable illness, and may have actually been a measurement of early in vivo activity of the CAR-T cells^82–84^. The animals exhibited no neurotoxic symptoms or signs of fever and weight loss, which were described in a recent study of neurotoxicity following CAR-T cell infusion^85^. Previous rhesus macaque adoptive transfer studies found that a significant fraction of the infused T cells localize to the lung with limited to undetectable persistence in blood or lymphoid tissues^86,87^. In lung tissues, we detected very low levels of CTV-CAR/CXCR5-T cells in vivo from the animal sacrificed at 2 DPT. We detected CAR/CXCR5-T cells in BAL fluid in four of the six treated animals between 2 and 14 DPT; however, this accumulation was transient as no cells were detected at 28 DPT.

We have previously shown that CAR/CXCR5-T cells produced from chronically infected animals, while effective at suppressing SIV in culture, retain residual virus^70^. T1 animals received CAR/CXCR5-T cells generated using PBMCs from chronically SIV-infected animals. Thus, it was not surprising that T1 animals showed a spike in viral load immediately post-infusion. Nonetheless, even with the infusion of virus along with the CAR/CXCR5-T cells, one of the three treated animals maintained viral load below 10^4^ copies/mL throughout the study, and two had undetectable viral loads at necropsy.

While our animal numbers per group were too small to perform statistical analyses between groups, we found that two of the three T2 animals maintained viral loads below 10^4^ copies/mL for the entire study, with levels below or near the level of detection for up to 10 months post-infusion. We detected very little to no SIV RNA^+^ cells in both F and EF areas of LNs from these two animals one-month post-treatment. Since the CAR/CXCR5-T cells did not persist beyond one month, it is unclear how the animals maintained control for the subsequent months.

However, it is rare for rhesus macaques to spontaneously control viral rebound after interruption of ART ^88,89^. The CAR/CXCR5-T cells may have effectively suppressed and slowed down SIV recrudescence post-ART release, allowing the endogenous immune response to be effective in maintaining low or undetectable viral loads in these animals. The T2 animal that did not control viral infection showed several important differences from the two controller animals. In the non-controlling animal, the infused cells showed decreased expansion in vivo, declined more quickly, and were more abundant in the BAL. Additionally, the non-controlling animal had a relatively lower percentage central memory cells and lower CCR7 expression than the animals that controlled plasma viremia post-infusion. It is possible that, due to differences in central memory phenotype, the cells did not persist as long, or were not present at a sufficient levels, at the time of viral recrudescence, and failed to exert an effect on viral load.

CAR/CXCR5-T cells accumulated predominantly in lymphoid tissues, where they reached peak accumulation at 6 DPT and persisted for up to 28 days. The preferential homing of infused CAR/CXCR5-T cells to lymphoid tissues in our study may be due to relatively high frequency of central memory cells (median 63%)^90^. Overall, no CAR/CXCR5-T cells were found in the blood or tissues past 28 DPT. Many adoptive transfer studies utilize a lymphodepletion regimen using cytotoxic agents, such as Cytoxan, to create room for the adoptively transferred cells to implant^91–94^, and such a conditioning regimen in future studies may allow greater expansion and persistence of CAR/CXCR5-T cells. Alternatively, persistence might be improved by the addition of antigen-expressing cells at, or shortly after, the time of CAR-T cell infusion to stimulate CAR-T cells in vivo, as was recently reported by Rust et al^74^.

To achieve better long-term control of infection, CAR/CXCR5-T cell immunotherapy could also be combined with other strategies aimed at eliminating the HIV/SIV reservoir. Such therapies might include latency reversal agents^62,95^ or agents that modify the immune system^96^, such as an IL-15 superagonist^97^.

These studies provide preliminary evidence of the safety and efficacy of CAR/CXCR5-T cells in a rhesus macaque model of HIV. Future studies with larger numbers of animals are essential to definitively determine the safety and efficacy of this intervention. With their ability to localize to the site of viral replication and to interact with virally infected cells, these immunotherapeutic cells have the potential to play an important role in the long-term cure of HIV infection without the use of life-long ART.

## Materials and Methods

### Animal Study Design

The studies used 10 rhesus macaques that were positive for the class I allele *Mamu-A1*001* but negative for *Mamu-B*008* and *Mamu-B*017:01.* Rhesus macaques were housed at the Wisconsin National Primate Research Center (WNPRC). All procedures were approved by the University of Wisconsin-Madison College of Letters and Sciences and Vice Chancellor for Research and Graduate Education Centers Institutional Animal Care and Use Committee (IACUC protocol number G005529). The animal facilities of the Wisconsin National Primate Research Center are licensed by the US Department of Agriculture and accredited by AAALAC. Animals were monitored twice daily by veterinarians for any signs of disease, injury, or psychological abnormalities. At the conclusion of the study, animals were humanely euthanized by anesthetizing with ketamine (at least 15 mg/kg IM) or other form of WNPRC veterinary approved general anesthesia followed by an IV overdose (at least 50 mg/kg or to effect) of sodium pentobarbital or equivalent as approved by a WNPRC veterinarian. Death was defined by stoppage of the heart as determined by a qualified and experienced person using a stethoscope to monitor heart sounds from the chest area, as well as all other vital signs, which can be monitored by observation.

The initial treated animal was chronically infected with SIVmac239 for 20 months prior to treatment. The animal was necropsied at day 2 post-infusion to determine the abundance and localization of the infused cells. Pilot study 1 and 2 treated animals (T1 and T2) and control untreated animals (C) (n=3 per group) were infected intrarectally with SIVmac251 (1 × 10^8^ viral RNA). ART consisting of 5.1 mg/kg Tenofovir Disoproxil Fumerate (TDF) (Gilead), 40 mg/kg Emtricitabine (FTC) (Gilead) and 2.5 mg/kg Dolutegravir (DTG) (Viiv) was formulated at Beth Israel Deaconess Medical Center (BIDMC). ART was initiated at day 63–68 post-infection and continued daily until the day of cell infusion. Blood samples were drawn biweekly to monitor viral loads, and all animals had undetectable viral loads at the time of infusion. Animals were ART-suppressed for times indicated in Table 1. PBMCs were collected by density gradient centrifugation from blood draws either post-infection for T1 or pre-infection for T2. PBMCs were cryopreserved in CryoStor CS5 (BioLife Solutions Inc.) at a concentration between 4 and 20 million cells/mL and transported and stored in liquid nitrogen until use.

### Cell manufacturing and infusion

The CD4-MBL CAR/ CXCR5 construct was described previously^61^. The bi-specific CAR contains rhesus CD4 and MBL domains, which leads to specificity for SIV, linked to extracellular hinge, transmembrane and co-stimulatory domains of rhesus CD28^12,23^ . The follicular homing receptor, CXCR5, is linked to the CAR with a self-cleaving peptide, P2A. Gammaretroviruses were produced by lipofectamine-mediated transfection of 293T cells^61^. CD4-MBL CAR/CXCR5-T cells were manufactured using the CD4-MBL CAR/CXCR5 gammaretrovirus as outlined previously^70,98^. Prior to infusion, cells were stained with CTV, an intracellular fluorescent dye, resuspended at a density of 2 × 10^7^ cells/mL in RPMI for T0 and in PBS containing 10% autologous serum for all other animals, packed on ice and transported to the WNPRC. The CAR/CXCR5-T cells were infused intravenously over 20 min while the animals were sedated. A veterinarian was present during the entire infusion. The dose of cells ranged from 0.35 to 2 × 10^8^ cells/kg (Table 2). Following infusion, animals were evaluated for signs of pain, illness, and stress observing appetite, stool, typical behavior, and physical condition by the staff of the Animal Services Unit at least twice daily. The weight of the animals was monitored routinely throughout the protocol.

### Tissue, blood, and cell collection

Blood samples were drawn for viral load determination immediately before and after infusion and on days 2, 6, 10, 14 and then biweekly until necropsy. Complete blood counts (CBC) were monitored biweekly throughout the experiment. LN biopsies and BAL samples were collected on days 2, 6, 14, 28 and 60–69 post-infusion. Colon and rectal biopsies were collected on days 2, 14, 28 and 60–69 post-infusion. Animals were necropsied between day 69 and 302 post-infusion.

### Viral load determination

Viral loads were measured by Virology Services (WNPRC). vRNA was isolated from plasma samples using the Maxwell Viral Total Nucleic Acid Purification kit on the Maxwell 48RSC instrument (Promega, Madison WI). vRNA was then quantified using a highly sensitive qRT-PCR assay based on the one developed by Cline et al.^99^. RNA was reverse transcribed and amplified using TaqMan Fast Virus 1-Step Master Mix qRT-PCR Master Mix (Invitrogen) on the LightCycler 480 or LC96 instrument (Roche, Indianapolis, IN) and quantified by interpolation onto a standard curve made up of serial ten-fold dilutions of in vitro transcribed RNA. RNA for this standard curve was transcribed from the p239gag_Lifson plasmid, kindly provided by Dr. Jeffrey Lifson, (NCI/Leidos). The final reaction mixtures contained 150 ng random primers (Promega, Madison, WI), 600 nM each primer and 100 nM probe. Primer and probe sequences are as follows: forward primer: 5′-GTCTGCGTCATCTGGTGCATTC-3′, reverse primer: 5′-CACTAGCTGTCTCTGCACTATGTGTTTTG-3′ and probe: 5′-6-carboxyfluorescein-CTTCCTCAGTGTGTTTCACTTTCTCTTCTGCG-BHQ1-3′. The reactions cycled with the following conditions: 50°C for 5 min, 95°C for 20 s followed by 50 cycles of 95°C for 15 s and 62°C for 1 min. The limit of detection of this assay is 100 copies/mL.

### Flow cytometry

Multiparametric flow cytometry was performed on fresh, transduced PBMCs and on thawed PBMCs or BAL cells collected post-infusion with monoclonal antibodies cross-reactive in rhesus macaques to detect (CD4-MBL)CAR/CXCR5-T cells and SIV-specific T cells. Cells were incubated with Live/Dead NIR (Invitrogen); Alexa Fluor 700 mouse anti-human CD3 (SP34-2), FITC, Brilliant Violet 650 mouse anti-human CD4 (M-T477), Brilliant Violet 510 mouse anti-human CD8 (RPA-T8), PerCP/Cy5.5 mouse anti-human CD95 (DX2) and Brilliant Violet 605 mouse anti-human CD28 (28.2) (BD Biosciences); Phycoerythrin (PE) mouse anti-human CXCR5 (MU5UBEE) (eBiosciences); MBL (3E7) (Invitrogen) conjugated to Alexa Fluor 647. To detect SIV-specific CD8^+^ T cells, samples were incubated with PE-labeled GAG-CM9 (NIH Tetramer Core) at 37°C for 15 min.

### CAR/CXCR5-T cell PCR

DNA qPCR was used to determine the quantity of CAR-T cells in PBMCs and BAL. Genomic DNA was isolated from freshly thawed PBMCs or BAL collected post-infusion using the DNeasy Blood and Tissue kit (Qiagen). PCR primers were designed to specifically bind to the junction of the CD4 and MBL fragments of the CAR to avoid recognition of endogenous CD4 or MBL. The assay included 300 nM concentrations of the following primers: CAR forward primer 5′-ATATTGTGGTCCTGGCCTTTCA-3′; CAR reverse primer 5′-AAGAATTTGTTTCCGACCTGCC-3′; albumin forward primer 5′-TGCATGAGAAAACGCCAGTAA-3′; albumin reverse primer 5′-ATGGTCGCCTGTTCACCAA-3′. PCR was run on a CFX96 thermal cycler (BioRad) with a program of one cycle of denaturation at 95℃ for 2 min, followed by 40 cycles of 95℃ for 10 s and 60℃ for 30 s. An amplified DNA fragment of the CAR was used in a standard curve to determine the copy number of the CAR. Albumin, which is present as two copies per cell, was used to determine cell number. The limit of detection was 2 copies of CAR DNA per 10^5^ cells.

### Luminex assay

Serum samples were stored at -80°C prior to analysis. Samples were tested by the Cytokine Reference Laboratory (University of Minnesota) using the magnetic bead set PRCYTOMAG-40K (EMD Millipore). Samples were analyzed for Non-Human Primate (NHP)-specific TNFα, IFNγ, IL-6 & IL-2 using the Luminex platform and performed as a multi-plex. Fluorescent color-coded beads coated with a specific capture antibody were added to each sample. After incubation and washing, biotinylated detection antibody was added followed by phycoerythrin-conjugated streptavidin. The beads were read on a Luminex instrument (Bioplex 200). Samples were run in duplicate and values were interpolated from five-parameter fitted standard curves.

### Singleplex RNAScope in situ hybridization and immunohistochemistry

RNAScope in situ hybridization utilized the 2.5 HD Reagent RED kit (Advanced Cell Diagnostics) as described previously^65,66,100^ with modifications. FFPE tissue sections (5 µm) on slides were deparaffinized by baking 1 h at 60℃, rinsing in xylene followed by absolute ethanol and air-drying. Sections were boiled in RNAScope® 1× Target Retrieval buffer (Advanced Cell Diagnostics) for epitope retrieval. Sections were then washed in dH_2_O, dipped in absolute ethanol and air-dried. Following a protease pretreatment, sections were rinsed with dH_2_O and hybridized overnight at 40°C with one of the following probes (all from Advanced Cell Diagnostics): SIVmac239 no-env antisense probe or a custom-made probe for the gammaretroviral vector to detect the CAR/CXCR5-transduced cells, DapB probe as a negative control probe or Macaca mulatta peptidylprolyl isomerase B (cyclophilin B) probe served as a positive control. Sections were then washed with 0.5× RNAScope wash buffer (Advanced Cell Diagnostics) and incubated with amplification reagents (1-6) according to the manufacturer’s instructions. For chromogenic detection, sections were incubated with 120 µL of fast Red chromagen solution and washed as recommended by the manufacturer. For immunofluorescence staining, sections were blocked with 4% normal goat serum (NGS) and incubated overnight with the following primary antibodies: mouse-anti-human CD20 (Clone L26, Biocare), mouse-anti-CD68 (KP1; Biocare), rabbit anti-CD20 (Polyclonal, Thermo Scientific), rabbit anti-CD3 (SP7; Labvision/Thermo Scientific), rabbit anti-CD4 (EPR6855; Abcam), or rabbit anti-Ki67 (Clone SP6, Invitrogen/Thermo Scientific). Goat secondary antibodies (Jackson Immunoresearch Laboratories) against mouse, rabbit, or human IgM conjugated to Alexa Fluor^TM^ 488, Alexa Fluor^TM^ 647 or Cy5 were used. Sections were counterstained with 1µg/mL DAPI and mounted in Prolong® Gold (ThermoFisher Scientific).

### Duplex in situ hybridization combined with immunofluorescence

For simultaneous visualization of both SIV vRNA and CAR/CXCR5-transduced cells, the RNAScope multiplex fluorescent kit V2 (Advanced Cell Diagnostics) was used with the opal fluorophores system (Akoya Bioscience) according to the manufacturer’s instructions and as described previously^66^ with some modifications. In brief, 5µm FFPE tissue sections on slides were deparaffinized as described above. Sections were pretreated with H_2_O_2_ (to block endogenous peroxidase activity) and washed in dH_2_O. Heat-induced epitope retrieval was achieved by boiling sections in RNAScope® 1× target retrieval buffer (Advanced Cell Diagnostics). Sections were washed, dehydrated in absolute ethanol, and air-dried. Sections were incubated with protease solution, rinsed twice in dH_2_O and incubated with pre-warmed premixed target probes (all from Advanced Cell Diagnostics) in which SIVmac239 no env antisense probe channel 2 (C2) was diluted in the custom-made probe for the gammaretroviral vector to detect the CAR/CXCR5-transduced cells channel 1 (C1) at C2: C1 1:50 ratio overnight at 40°C. Sections were washed with a 0.5× RNAScope wash buffer. Amplification and HRP-C1 and HRP-C2 signal development were performed as recommended by the manufacturer with the modification of the use of 0.5× RNAScope wash buffer instead of 1× RNAScope wash buffer and use of a 1:150 dilution of Opals (all from Akoya Bioscience) instead of 1:1500. Opal™ 570 and Opal™ 690 were used for C1 and C2, respectively. For immunofluorescence staining, sections were washed twice in TBST (TBS-tween 20–0.05% v/v), blocked in 10% NGS-TBS-1% BSA and incubated with primary antibodies diluted in TBS-1% BSA for 1 h at RT. Primary antibodies included the same antibodies described in Singleplex vRNA in situ hybridization combined with immunofluorescence. The sections were washed and incubated with secondary antibodies, Opal Polymer HRP Ms + Rb for 10 min at RT. After washing, the sections were incubated with Opal™ 520 diluted 1:150 in the multiplex TSA buffer (Advanced Cell Diagnostics) for 10 min at RT. After washing, sections were counterstained with 1µg/mL DAPI and mounted in Prolong® Gold (ThermoFisher Scientific).

### Quantitative image analysis for RNAScope

Sections were imaged using a Leica DM6000 confocal microscope. Montage images of multiple 512 × 512 pixels were created and used for analysis. F and EF areas were delineated using Leica software with B cell follicle areas identified morphologically as clusters of closely aggregated brightly stained CD20^+^ or IgM^+^ cells. Some sections were co-stained with goat anti-human IgM-AF647 (Jackson ImmunoResearch) and mouse-anti-human CD20 antibodies (clone L26, Biocare Medical, Inc.) to confirm that both antibodies co-localized similarly in B cell follicles. Cell counts were done using LAS X (Leica confocal) software; each cell was demarcated using a Leica software tool to avoid counting the same cell twice. Leica software was used to measure the delineated areas for cell counts. To determine the percentage of follicles that had CAR/CXCR5-T cells over time post-infusion, a total of 790 follicles were evaluated for presence of CAR/CXCR5-T cells with a median of 302 follicles (range, 172–316) per animal. To determine the levels of CAR/CXCR5-T cells/mm^2^ in follicular areas, over time post-infusion, a median of 8.4 mm^2^ (range, 6.8–8.9 mm^2^) of follicular area was analyzed with a total of 172 follicles analyzed with a median of 57 follicles (range, 48–67) per animal. In addition, a total of 190 follicles were examined to determine the percentage of follicles that had a cluster of expanding CAR/CXCR5-T cells at the edge of the follicle at 2 DPT, with a median of 95 follicles (range 90–100). To determine the percentage of follicles with free virions bound by FDC over time post-infusion, a total of 518 follicles with a median of 146 follicles (range 140– 232) per animal. To determine the levels of SIV RNA^+^ cells/mm^2^ in follicular areas, over time post-infusion, a median of 6.46 mm^2^ (range 5.48–6.84 mm^2^) of follicular area was analyzed with a total of 131 follicles analyzed with a median of 45 follicles (range 38–48) per animal. To determine levels of CAR/CXCR5-T cells/mm^2^ and level of SIV RNA^+^ cells/mm^2^ in treated animals, a median of 19.9 mm^2^ (range of 15.7–34 mm^2^) of EF areas were analyzed per animal. To confirm the specificity of the custom-made probe that we designed to detect the gammaretroviral CAR/CXCR5 construct and to determine the level of SIV vRNA^+^ cells in F areas, LN tissues from three untreated control animals were hybridized to the custom made probe and an SIV probe; a median of 2.36 mm^2^ (range 0.79–2.99 mm^2^) of follicular area was analyzed with a total of 53 follicles analyzed with a median of 22 follicles (range 5–26) per animal. To determine the level of SIV RNA^+^ cells/mm^2^ in untreated control animals, a median of 3.6 mm^2^ (range of 0.68–9.3 mm^2^) of EF areas was analyzed per animal.

### Immunohistochemistry and analyses

Indirect immunohistochemistry was performed on fresh tissue specimens shipped overnight, sectioned with a compresstome^101^ and stained as described previously^102–104^. Briefly, sections were stained with 0.4 µg/mL rabbit -anti-human CD20 polyclonal antibodies (Neomarkers) and 2 µg/mL rat-anti-human CD3 antibodies (clone MCA1477, BioRad). Then, sections were stained with secondary antibodies by incubating with 0.3 µg/mL Alexa Fluor^TM^ 488-conjugated goat-anti-rabbit antibodies, and 0.2–0.3 µg/mL Cy5-conjugated goat anti-rat antibodies overnight at 4°C. Secondary antibodies were obtained from Jackson ImmunoResearch Laboratories (West Grove, PA). Sections were imaged using a Leica DM6000 confocal microscope. Montage images of multiple 512 × 512 pixels were created and used for analysis. Confocal z-series were collected in a step size of 3 µm. Images were opened and analyzed in LAS X (Leica confocal) software directly. We used the LAS X software to create montages of multiple projected confocal serial z-scans. Follicular areas were identified morphologically as clusters of brightly stained, closely aggregated CD20^+^ cells. F and EF areas were delineated and measured using LAS X software. Areas were not included if they showed loosely aggregated B cells that were ambiguous. To prevent bias, the yellow CTV channel was turned off when F and EF areas were delineated. Cell counts were performed on single z-scans.

### Statistical analyses

Statistical analyses utilized GraphPad Prism 8.3.0 for Windows (GraphPad Software, San Diego, CA). Specific tests are indicated in the figure legends. Correlations were determined using Spearman’s correlation, assuming independence.

## Acknowledgments

Anti-CD3 and anti-CD28 used in these studies were provided by the NIH Nonhuman Primate Reagent Resource (R24 OD010976, U24 AI126683). IL-2 used in these studies was provided by The NCI Preclinical Repository. GAG-CM9 tetramers were provided by the NIH Tetramer Core.

The authors thank the following University of Minnesota scientists: Ms. Preethi Haran for tissue staining, Ms. Chi Phan for virus preparation, Ms. Jodi Anderson, Mr. Steve Wietgrefe and Dr. Lijie Duan for assistance with RNAScope development, Mr. Michael Ehrhardt of the Cytokine Reference Laboratory for the Luminex assay; the following staff members at the WNPRC: Dr. Heather Simmons for pathology services, Mr. Dane Schalk for animal services, Dr. Nancy Schultz-Darken for animal project oversight; the following staff members at the University of Wisconsin, Madison: Ms. Kim Weisgrau for cell isolation and flow cytometry, Ms. Andrea Weiler for viral load determination. We also thank Dr. Jacob Estes and Dr. Kathleen Busman-Sahay of Oregon Health and Science University for assistance with RNAScope development, Dr. Mauricio Martins at the University of Miami for the donation of the chronically infected animal to the WNPRC and Dr. James Whitney of Harvard University for MHC genotyping the animals, and for the viral load data from the C and T1 group animals during early infection and ART prior to these animals being provided to our study. We also thank Natalie Coleman Fuller for assistance in editing this manuscript.

## Author contributions

**Conceptualization:** Pamela Skinner

**Data curation:** Mary Pampusch, Hadia Abdelaal

**Formal analysis:** Mary Pampusch, Hadia Abdelaal, Emily Cartwright, Aaron Rendahl

**Funding acquisition:** Eva Rakasz, Elizabeth Connick, Edward Berger, Pamela Skinner

**Investigation:** Mary Pampusch, Hadia Abdelaal, Emily Cartwright, Jhomary Molden, Brianna Davey, Jordan Sauve

**Methodology:** Mary Pampusch, Hadia Abdelaal

**Project Administration:** Mary Pampusch, Pamela Skinner

**Resources:** Eva Rakasz, Edward Berger, Pamela Skinner

**Supervision:** Eva Rakasz, Elizabeth Connick, Edward Berger, Pamela Skinner

**Visualization:** Mary Pampusch, Hadia Abdelaal, Emily Cartwright, Jhomary Molden, Brianna Davey

**Writing-original draft:** Mary Pampusch, Hadia Abdelaal, Pamela Skinner

**Writing-review and editing:** Emily Cartwright, Eva Rakasz, Elizabeth Connick, Edward Berger

## Supporting information

**Fig S1. Immunotherapeutic cell infusion leads to transient increases in cytokine levels and cell accumulation in bronchoalveolar lavage fluid.**

Serum samples were analyzed for post-infusion production of cytokines using a non-human primate (NHP)-specific (a) tumor necrosis factor (TNF) alpha, (b) interferon (IFN) gamma, (c) interleukin (IL)-6 and (d) IL-2 multiplex Luminex assay. Lung accumulation of CAR T cells was determined by analysis of bronchoalveolar lavage (BAL) samples. Cells were isolated from BAL and analyzed for (e) the percentage of CD4-MBL CAR^+^ cells in the CD3^+^ T population by flow cytometry or (f) the number of copies of CAR/cell in the total cell population by quantitative real-time PCR.

**Fig S2. Representative flow plots from cells prepared for infusion.** 1 × 10^6^ cells were stained with the antibodies listed in the Flow Cytometry section of Materials and Methods. Gating strategy for determination of co-expression of CAR (MBL) and CXCR5. The CD8+ population was used to determine central memory phenotype (CD28^+^CD95^+^) and CCR7 expression. Plots presented are from cells infused into Rh2858.

**Fig S3. Levels of tetramer^+^ CD3 T cells present in peripheral blood mononuclear cells (PBMCs)**. PBMC, collected on d28 from animals in each of the three groups, were stained for Gag CM9 and analyzed by flow cytometry as described in Materials and Methods. The bar represents the median.

**Fig S4. Numbers of follicular CAR/CXCR5-T cells identified in situ in lymph nodes correlate with numbers of CAR-specific PBMCs identified by flow cytometry.**

Correlation between follicular CAR/CXCR5-T cells/mm^2^ by RNAScope and CD4-MBL^+^ PBMC by flow cytometry. Association was tested using Spearman’s correlation. Scales are log (value+1) on the y-axis and log (value) on the x-axis; labels use the original units. The line represents the fitted regression.

## Notes

### Competing Interest Statement

Pamela Skinner is the co-founder and CSO of MarPam Pharma and has a patent pending US20180371057A1. Mary Pampusch was a former employee of MarPam Pharma. Other authors have no competing interests.

## References

1. WHO | HIV/AIDS. WHO (2020).

2. Zhang, L. et al. Quantifying Residual HIV-1 Replication in Patients Receiving Combination Antiretroviral Therapy. N. Engl. J. Med. 340, 1605–1613 (1999).

3. Wong, J. K. et al. Recovery of replication-competent HIV despite prolonged suppression of plasma viremia. Science (80-.). 278, 1291–1295 (1997).

4. Deeks, S. G. et al. A phase II randomized study of HIV-specific T-cell gene therapy in subjects with undetectable plasma viremia on combination antiretroviral therapy. Mol. Ther. 5, 788–797 (2002).

5. Fernandez-Montero, J. V., Eugenia, E., Barreiro, P., Labarga, P. & Soriano, V. Antiretroviral drug-related toxicities-clinical spectrum, prevention, and management. Expert Opinion on Drug Safety vol. 12 697–707 (2013).

6. Iacob, S. A., Iacob, D. G. & Jugulete, G. Improving the Adherence to Antiretroviral Therapy, a Difficult but Essential Task for a Successful HIV Treatment—Clinical Points of View and Practical Considerations. Front. Pharmacol. 8, 831 (2017).

7. Johnson Lyons, S., et al. Monitoring Selected National HIV Prevention and Care Objectives by Using HIV Surveillance Data—United States and 6 Dependent Areas, 2019. HIV Surveill. Suppl. Rep. 26,.

8. Montessori, V., Press, N., Harris, M., Akagi, L. & Montaner, J. S. G. Adverse effects of antiretroviral therapy for HIV infection. CMAJ 170, 229–38 (2004).

9. Reust, C. E. Common adverse effects of antiretroviral therapy for HIV disease. Am. Fam. Physician 83, 1443–1451 (2011).

10. WHO | HIV drug resistance. WHO (2019).

11. Ndung’u, T., McCune, J. M. & Deeks, S. G. Why and where an HIV cure is needed and how it might be achieved. Nature vol. 576 397–405 (2019).

12. Liu, L. et al. Novel CD4-Based Bispecific Chimeric Antigen Receptor Designed for Enhanced Anti-HIV Potency and Absence of HIV Entry Receptor Activity. J. Virol. 89, 6685–6694 (2015).

13. Katlama, C. et al. Barriers to a cure for HIV: New ways to target and eradicate HIV-1 reservoirs. The Lancet vol. 381 2109–2117 (2013).

14. Siliciano, J. D. & Siliciano, R. F. Recent developments in the search for a cure for HIV-1 infection: Targeting the latent reservoir for HIV-1. Journal of Allergy and Clinical Immunology vol. 134 12–19 (2014).

15. Lewin, S. R., Deeks, S. G. & Barré-Sinoussi, F. Towards a cure for HIV-are we making progress? The Lancet vol. 384 209–211 (2014).

16. Archin, N. M. & Margolis, D. M. Emerging strategies to deplete the HIV reservoir. Current Opinion in Infectious Diseases vol. 27 29–35 (2014).

17. Ananworanich, J. & Fauci, A. S. HIV cure research: a formidable challenge. J. virus Erad. 1, 1–3 (2015).

18. Neelapu, S. S. et al. Axicabtagene Ciloleucel CAR T-Cell Therapy in Refractory Large B-Cell Lymphoma. N. Engl. J. Med. 377, 2531–2544 (2017).

19. Maude, S. L. et al. Tisagenlecleucel in Children and Young Adults with B-Cell Lymphoblastic Leukemia. N. Engl. J. Med. 378, 439–448 (2018).

20. Schuster, S. J. et al. Tisagenlecleucel in Adult Relapsed or Refractory Diffuse Large B-Cell Lymphoma. N. Engl. J. Med. 380, 45–56 (2019).

21. Li, C., Mei, H. & Hu, Y. Applications and explorations of CRISPR/Cas9 in CAR T-cell therapy. Brief. Funct. Genomics 19, 175–182 (2020).

22. Depil, S., Duchateau, P., Grupp, S. A., Mufti, G. & Poirot, L. ‘Off-the-shelf’ allogeneic CAR T cells: development and challenges. Nat. Rev. Drug Discov. 19, 185–199 (2020).

23. Ghanem, M. H. et al. Bispecific chimeric antigen receptors targeting the CD4 binding site and high-mannose Glycans of gp120 optimized for anti–human immunodeficiency virus potency and breadth with minimal immunogenicity. Cytotherapy 20, 407–419 (2018).

24. Folkvord, J. M., Armon, C. & Connick, E. Lymphoid Follicles Are Sites of Heightened Human Immunodeficiency Virus Type 1 (HIV-1) Replication and Reduced Antiretroviral Effector Mechanisms. AIDS Res. Hum. Retroviruses 21, 363–370 (2005).

25. Hufert, F. T. et al. Germinal centre CD4+ T cells are an important site of HIV replication in vivo. AIDS 11, 849–57 (1997).

26. Connick, E. et al. CTL Fail to Accumulate at Sites of HIV-1 Replication in Lymphoid Tissue. J. Immunol. 178, 6975–6983 (2007).

27. Connick, E. et al. Compartmentalization of Simian Immunodeficiency Virus Replication within Secondary Lymphoid Tissues of Rhesus Macaques Is Linked to Disease Stage and Inversely Related to Localization of Virus-Specific CTL. J. Immunol. 193, 5613–5625 (2014).

28. Tenner-Racz, K. et al. The unenlarged lymph nodes of HIV-1-infected, asymptomatic patients with high CD4 T cell counts are sites for virus replication and CD4 T cell proliferation. The impact of highly active antiretroviral therapy. J. Exp. Med. 187, 949– 959 (1998).

29. Biberfeld, P. et al. HTLV-III expression in infected lymph nodes and relevance to pathogenesis of lymphadenopathy. Am. J. Pathol. 125, 436–442 (1986).

30. Brenchley, J. M. et al. Differential infection patterns of CD4+ T cells and lymphoid tissue viral burden distinguish progressive and nonprogressive lentiviral infections. Blood 120, 4172–81 (2012).

31. Perreau, M. et al. Follicular helper T cells serve as the major CD4 T cell compartment for HIV-1 infection, replication, and production. J. Exp. Med. 210, 143–56 (2013).

32. Fukazawa, Y. et al. B cell follicle sanctuary permits persistent productive simian immunodeficiency virus infection in elite controllers. Nat. Med. 21, 132–9 (2015).

33. Pope, M. & Haase, A. T. Transmission, acute HIV-1 infection and the quest for strategies to prevent infection. Nature Medicine vol. 9 847–852 (2003).

34. Pantaleo, G. et al. HIV infection is active and progressive in lymphoid tissue during the clinically latent stage of disease. Nature 362, 355–8 (1993).

35. Embretson, J. et al. Massive covert infection of helper T lymphocytes and macrophages by HIV during the incubation period of AIDS. Nature 362, 359–362 (1993).

36. Schacker, T. et al. Rapid Accumulation of Human Immunodeficiency Virus (HIV) in Lymphatic Tissue Reservoirs during Acute and Early HIV Infection: Implications for Timing of Antiretroviral Therapy. J. Infect. Dis. 181, 354–357 (2000).

37. Haase, A. T. et al. Quantitative image analysis of HIV-1 infection in lymphoid tissue. Science 274, 985–9 (1996).

38. Fox, C. H. et al. Lymphoid germinal centers are reservoirs of human immunodeficiency virus type 1 RNA. J. Infect. Dis. 164, 1051–1057 (1991).

39. Heath, S. L., Tew, J. G., Szakal, A. K. & Burton, G. F. Follicular dendritic cells and human immunodeficiency virus infectivity. Nature vol. 377 740–4 (1995).

40. Smith-Franklin, B. A. et al. Follicular Dendritic Cells and the Persistence of HIV Infectivity: The Role of Antibodies and Fcγ Receptors. J. Immunol. 168, 2408–2414 (2002).

41. Joling, P. et al. Binding of human immunodeficiency virus type-1 to follicular dendritic cells in vitro is complement dependent. J. Immunol. 150, (1993).

42. Tjernlund, A. et al. In situ detection of Gag-specific CD8+ cells in the GI tract of SIV infected Rhesus macaques. Retrovirology 7, 12 (2010).

43. Sasikala-Appukuttan, A. K. et al. Location and Dynamics of the Immunodominant CD8 T Cell Response to SIVΔnef Immunization and SIVmac251 Vaginal Challenge. PLoS One 8, e81623 (2013).

44. Li, S. et al. Low levels of siv-specific CD8+ T cells in germinal centers characterizes acute SIV infection. PLoS Pathog. 15, (2019).

45. Li, Q. et al. Visualizing antigen-specific and infected cells in situ predicts outcomes in early viral infection. Science 323, 1726–9 (2009).

46. Li, S. et al. Simian Immunodeficiency Virus-Producing Cells in Follicles Are Partially Suppressed by CD8+ Cells In Vivo. J. Virol. 90, 11168–11180 (2016).

47. Förster, R. et al. A Putative Chemokine Receptor, BLR1, Directs B Cell Migration to Defined Lymphoid Organs and Specific Anatomic Compartments of the Spleen. Cell 87, 1037–1047 (1996).

48. Schaerli, P. et al. CXC chemokine receptor 5 expression defines follicular homing T cells with B cell helper function. J. Exp. Med. 192, 1553–62 (2000).

49. Haynes, N. M. et al. Role of CXCR5 and CCR7 in follicular Th cell positioning and appearance of a programmed cell death gene-1high germinal center-associated subpopulation. J. Immunol. 179, 5099–108 (2007).

50. Quigley, M. F., Gonzalez, V. D., Granath, A., Andersson, J. & Sandberg, J. K. CXCR5+ CCR7- CD8 T cells are early effector memory cells that infiltrate tonsil B cell follicles. Eur. J. Immunol. 37, 3352–3362 (2007).

51. Gunn, M. D. et al. A B-cell-homing chemokine made in lymphoid follicles activates Burkitt’s lymphoma receptor-1. Nature 391, 799–803 (1998).

52. Legler, D. F. et al. B cell-attracting chemokine 1, a human CXC chemokine expressed in lymphoid tissues, selectively attracts B lymphocytes via BLR1/CXCR5. J. Exp. Med. 187, 655–660 (1998).

53. Havenar-Daughton, C. et al. CXCL13 is a plasma biomarker of germinal center activity. Proc. Natl. Acad. Sci. U. S. A. 113, 2702–2707 (2016).

54. Chang, J. E. & Turley, S. J. Stromal infrastructure of the lymph node and coordination of immunity. Trends in Immunology vol. 36 30–39 (2015).

55. Katakai, T. et al. Organizer-Like Reticular Stromal Cell Layer Common to Adult Secondary Lymphoid Organs. J. Immunol. 181, 6189–6200 (2008).

56. Katakai, T. Marginal reticular cells: a stromal subset directly descended from the lymphoid tissue organizer. Front. Immunol. 3, 200 (2012).

57. Wang, X. et al. Follicular dendritic cells help establish follicle identity and promote B cell retention in germinal centers. J. Exp. Med. 208, 2497–2510 (2011).

58. Kroenke, M. A. et al. Bcl6 and Maf Cooperate To Instruct Human Follicular Helper CD4 T Cell Differentiation. J. Immunol. 188, 3734–3744 (2012).

59. Rasheed, A.-U., Rahn, H.-P., Sallusto, F., Lipp, M. & Müller, G. Follicular B helper T cell activity is confined to CXCR5hiICOShi CD4 T cells and is independent of CD57 expression. Eur. J. Immunol. 36, 1892–1903 (2006).

60. Ayala, V. I. et al. CXCR5-Dependent Entry of CD8 T Cells into Rhesus Macaque B-Cell Follicles Achieved through T-Cell Engineering. J. Virol. 91, (2017).

61. Haran, K. P. et al. Simian Immunodeficiency Virus (SIV)-Specific Chimeric Antigen Receptor-T Cells Engineered to Target B Cell Follicles and Suppress SIV Replication. Front. Immunol. 9, 1–12 (2018).

62. Skinner, P. J. Targeting reservoirs of HIV replication in lymphoid follicles with cellular therapies to cure HIV. Adv. Cell Gene Ther. 2, e27 (2019).

63. Skinner, P. J. Overcoming the Immune Privilege of B cell Follicles to Cure HIV-1 Infection. 1, 1–3 (2014).

64. Wang, F. et al. RNAscope: A novel in situ RNA analysis platform for formalin-fixed, paraffin-embedded tissues. J. Mol. Diagnostics 14, 22–29 (2012).

65. Deleage, C. et al. Defining HIV and SIV Reservoirs in Lymphoid Tissues. Pathog. Immun. 1, 68 (2016).

66. Vasquez, J. J. et al. Elucidating the Burden of HIV in Tissues Using Multiplexed Immunofluorescence and In Situ Hybridization: Methods for the Single-Cell Phenotypic Characterization of Cells Harboring HIV In Situ. J. Histochem. Cytochem. 66, 427–446 (2018).

67. Ayala, V. I. et al. Adoptive Transfer of Engineered Rhesus Simian Immunodeficiency Virus-Specific CD8 + T Cells Reduces the Number of Transmitted/Founder Viruses Established in Rhesus Macaques . J. Virol. 90, 9942–9952 (2016).

68. Randolph, G. J. CCR7: Unifying Disparate Journeys to the Lymph Node. J. Immunol. 196, 3–4 (2016).

69. Förster, R. et al. CCR7 coordinates the primary immune response by establishing functional microenvironments in secondary lymphoid organs. Cell 99, 23–33 (1999).

70. Pampusch, M. S. et al. Rapid Transduction and Expansion of Transduced T Cells with Maintenance of Central Memory Populations. Mol. Ther. - Methods Clin. Dev. 16, 1–10 (2020).

71. Bronnimann, M. P., Skinner, P. J. & Connick, E. The B-cell follicle in HIV infection: Barrier to a cure. Front. Immunol. 9, 1–13 (2018).

72. Mylvaganam, G. H. et al. Dynamics of SIV-specific CXCR5+ CD8 T cells during chronic SIV infection. Proc. Natl. Acad. Sci. U. S. A. 114, (2017).

73. Li, S. et al. Simian immunodeficiency virus-producing cells in follicles are partially suppressed by CD8^+^ cells in vivo. J. Virol. 90, (2016).

74. He, R. et al. Follicular CXCR5-expressing CD8+ T cells curtail chronic viral infection. (2016) doi:10.1038/nature19317.

75. Younan, P. M. et al. Lentivirus-mediated gene transfer in hematopoietic stem cells is impaired in SHIV-infected, ART-treated nonhuman primates. Mol. Ther. 23, 943–951 (2015).

76. Ollerton, M. T., Berger, E. A., Connick, E. & Burton, G. F. HIV-1-Specific Chimeric Antigen Receptor T Cells Fail To Recognize and Eliminate the Follicular Dendritic Cell HIV Reservoir In Vitro. J. Virol. 94, (2020).

77. Bitton, N., Verrier, F., Debré, P. & Gorochov, G. Characterization of T cell-expressed chimeric receptors with antibody-type specificity for the CD4 binding site of HIV-1 gp120. Eur. J. Immunol. 28, 4177–4187 (1998).

78. Romeo, C. & Seed, B. Cellular immunity to HIV activated by CD4 fused to T cell or Fc receptor polypeptides. Cell 64, 1037–1046 (1991).

79. Sahu, G. K. et al. Anti-HIV designer T cells progressively eradicate a latently infected cell line by sequentially inducing HIV reactivation then killing the newly gp120-positive cells. Virology 446, 268–275 (2013).

80. Qi, J., Ding, C., Jiang, X. & Gao, Y. Advances in Developing CAR T-Cell Therapy for HIV Cure. Front. Immunol. 11, 1–13 (2020).

81. Brenchley, J. M. et al. CD4+ T cell depletion during all stages of HIV disease occurs predominantly in the gastrointestinal tract. J. Exp. Med. 200, 749–759 (2004).

82. Kalos, M. et al. T cells with chimeric antigen receptors have potent antitumor effects and can establish memory in patients with advanced leukemia. Sci. Transl. Med. 3, 95ra73 (2011).

83. Kalos, M. Biomarkers in T cell therapy clinical trials. Journal of Translational Medicine vol. 9 138 (2011).

84. Porter, D., Levine, B., Kalos, M., Bagg, A. & June, C. H. Chimeric Antigen Receptor– Modified T Cells in Chronic Lymphoid Leukemia. N. Engl. J. Med. 365, 725–733 (2011).

85. Taraseviciute, A. et al. Chimeric antigen receptor T cell–mediated neurotoxicity in nonhuman primates. Cancer Discov. 8, 750–763 (2018).

86. Bolton, D. L. et al. Trafficking, persistence, and activation state of adoptively transferred allogeneic and autologous Simian Immunodeficiency Virus-specific CD8(+) T cell clones during acute and chronic infection of rhesus macaques. J. Immunol. 184, 303–14 (2010).

87. Minang, J. T. et al. Distribution, persistence, and efficacy of adoptively transferred central and effector memory-derived autologous simian immunodeficiency virus-specific CD8+ T cell clones in rhesus macaques during acute infection. J. Immunol. 184, 315–326 (2010).

88. Strongin, Z. et al. Virologic and Immunologic Features of Simian Immunodeficiency Virus Control Post-ART Interruption in Rhesus Macaques. J. Virol. 94, (2020).

89. Borducchi, E. N. et al. Ad26/MVA therapeutic vaccination with TLR7 stimulation in SIV-infected rhesus monkeys. Nature 540, 284–287 (2016).

90. Ayala, V. I. et al. Adoptive Transfer of Engineered Rhesus Simian Immunodeficiency Virus-Specific CD8 T Cells Reduces the Number of Transmitted/ Founder Viruses Established in Rhesus Macaques. (2016) doi:10.1128/JVI.01522-16.

91. Anasetti, C. & Mulé, J. J. To ablate or not to ablate? HSCs in the T cell driver’s seat. Journal of Clinical Investigation vol. 117 306–310 (2007).

92. June, C. H. Adoptive T cell therapy for cancer in the clinic. Journal of Clinical Investigation vol. 117 1466–1476 (2007).

93. Greenberg, P. D. Adoptive T cell therapy of tumors: Mechanisms operative in the recognition and elimination of tumor cells. Adv. Immunol. 49, 281–355 (1991).

94. Dudley, M. E. et al. Cancer regression and autoimmunity in patients after clonal repopulation with antitumor lymphocytes. Science (80-.). 298, 850–854 (2002).

95. Sengupta, S. & Siliciano, R. F. Targeting the Latent Reservoir for HIV-1. Immunity vol. 48 872–895 (2018).

96. Mu, W., Carrillo, M. A. & Kitchen, S. G. Engineering CAR T Cells to Target the HIV Reservoir. Front. Cell. Infect. Microbiol. 10, 410 (2020).

97. Webb, G. M. et al. The human IL-15 superagonist ALT-803 directs SIV-specific CD8+ T cells into B-cell follicles. Blood Adv. 2, 76–84 (2018).

98. Pampusch, M. S. & Skinner, P. J. Transduction and expansion of primary T cells in nine days with maintenance of central memory phenotype. J. Vis. Exp. 2020, 60400 (2020).

99. Nichole Cline, A., Bess, J. W., Piatak, M. J. & Lifson, J. D. Highly sensitive SIV plasma viral load assay: practical considerations, realistic performance expectations, and application to reverse engineering of vaccines for AIDS. J. Med. Primatol. 34, 303–312 (2005).

100. Bertram, K. M. et al. Identification of HIV transmitting CD11c+ human epidermal dendritic cells. Nat. Commun. 10, 1–15 (2019).

101. Abdelaal, H. M. et al. Comparison of Vibratome and Compresstome sectioning of fresh primate lymphoid and genital tissues for in situ MHC-tetramer and immunofluorescence staining. Biol. Proced. Online 17, 2 (2015).

102. Skinner, P. J., Daniels, M. a., Schmidt, C. S., Jameson, S. C. & Haase, a. T. Cutting Edge: In Situ Tetramer Staining of Antigen-Specific T Cells in Tissues. J. Immunol. 165, 613– 617 (2000).

103. Li, S., Mwakalundwa, G. & Skinner, P. J. In situ MHC-tetramer staining and quantitative analysis to determine the location, abundance, and phenotype of antigen-specific CD8 T cells in tissues. J. Vis. Exp. 2017, 1–8 (2017).

104. Abdelaal, H. M., Cartwright, E. K. & Skinner, P. J. Detection of Antigen-Specific T Cells Using In Situ MHC Tetramer Staining. Int. J. Mol. Sci. 20, (2019).

